# Remodeling of the H3 nucleosomal landscape during mouse aging

**DOI:** 10.1101/769489

**Authors:** Yilin Chen, Juan I. Bravo, Bérénice A. Benayoun

**Affiliations:** Leonard Davis School of Gerontology, University of Southern California, Los Angeles, CA 90089, USA; Graduate program in the Biology of Aging, University of Southern California, Los Angeles, CA 90089, USA; USC Norris Comprehensive Cancer Center, Epigenetics and Gene Regulation, Los Angeles, CA 90089, USA; USC Stem Cell Initiative, Los Angeles, CA 90089, USA

## Abstract

In multi-cellular organisms, the control of gene expression is key not only for development, but also for adult cellular homeostasis, and deregulation of gene expression correlates with aging. A key layer in the study of gene regulation mechanisms lies at the level of chromatin: cellular chromatin states (*i.e.* the ‘epigenome’) can tune transcriptional profiles, and, in line with the prevalence of transcriptional alterations with aging, accumulating evidence suggests that the chromatin landscape is altered with aging across cell types and species. However, though alterations in the chromatin make-up of cells are considered to be a hallmark of aging, little is known of the genomic loci that are specifically affected by age-related chromatin state remodeling and of their biological significance. Here, we report the analysis of genome-wide profiles of core histone H3 occupancy in aging male mice tissues (i.e. heart, liver, cerebellum and olfactory bulb) and primary cultures of neural stem cells. We find that, although no drastic changes in H3 levels are observed, local changes in H3 occupancy occur with aging across tissues and cells with both regions of increased or decreased occupancy. These changes are compatible with a general increase in chromatin accessibility at pro-inflammatory genes and may thus mechanistically underlie known shift in gene expression programs with aging.

## 1. Introduction

Precise control of gene expression is key not only for development in metazoans, but also for adult cellular homeostasis, and deregulation of gene expression correlates with aging [1–3]. Aging is the main risk factor for many chronic diseases, including neurodegeneration, cardiovascular disease, diabetes, and cancer. In eukaryotes, transcriptional profiles are regulated in part by chromatin states, which are notably governed by post-translational modifications of core histone proteins [4–9]. Interestingly, remodeling of chromatin landscapes, or so-called “epigenetic alterations” is considered to be a hallmark of aging [10, 11].

The overwhelming majority of core histone expression is restricted to the S-phase of the cell cycle [12], and post-mitotic or terminally differentiated cells show very little *de novo* synthesis [13]. Indeed, a number of canonical and variant histones proteins are among the longest-lived proteins in the proteomes of rat brain and liver, with stability in the order of months [14, 15]. Interestingly, core histone protein levels decrease during yeast replicative aging [16], and in mammalian models of cellular senescence [17]. Both of these paradigms are the results of repeated accumulated cell divisions. In addition, aged mouse muscle stem cells have lower transcript levels of several histone genes [18]. Substantial changes in histone expression level could lead to genome-wide remodeling of nucleosomal occupancy, and global changes in transcriptional outputs. Indeed, decreased histone expression during yeast replicative aging is linked to an overall decrease in nucleosome occupancy and the aberrant gene upregulation [19], and experimental modulation of histone exchange and deposition into chromatin, or overexpression of histone H3 and H4, can modulate *S. cerevisiae* lifespan [16]. Interestingly, even substantial loss of histone protein levels may lead to nucleosome occupancy changes at only a small number of genomic loci [20], thus suggesting that nucleosome occupancy may represent an important layer of gene expression regulation.

Local changes in nucleosome occupancy have been observed in aging liver using MNase-seq [21], which may facilitate the age-related activation of lipogenesis genes [21]. However, whether the observed remodeling results from changes in core histone expression or deposition patterns is unclear. In addition, whether mammalian aging outside of highly replicative compartments is associated with histone loss throughout aging remains unknown.

Here, we present a reanalysis of H3 ChIP-seq datasets derived from a recent epigenomic study of male mouse aging across 4 tissues (i.e. heart, liver, cerebellum, olfactory bulb) and one primary cell type (*i.e.* primary neural stem cell cultures [NSCs]) at 3 ages throughout mouse lifespan (3, 12 and 29 months of age[22]. Although we do not find evidence for large changes in histone H3 expression levels or nucleosome occupancy across ages, we do identify reproducibly remodeled nucleosomes with mouse aging. This analysis represents an important resource in the study of epigenomic remodeling during mammalian aging.

## 2. Results

### 2.1. Analysis of H3 ChIP-seq datasets reveals limited remodeling of H3 nucleosomes with aging

We took advantage of our previously obtained H3 ChIP-seq samples from aging male mouse tissue and cell samples to identify differentially regulated H3 nucleosome regions with aging [22] (Fig. 1A). Consistent with a previous study using MNase-seq on mouse aging liver tissue [21], we found that significant histone occupancy changes with mouse chronological aging were restricted to a limited number of loci (Fig. 1B-D). We were able to identify regions with both increased H3 occupancy or decreased H3 occupancy with aging (Fig. 1B-D, **S1**), suggesting that a remodeling of accessible and inaccessible regions occurs during aging. Regions with increased H3 occupancy with aging are expected to correspond to regions that are less accessible to the transcriptional machinery in aged tissues. Conversely, regions with decreased H3 occupancy with aging are expected to correspond to regions that are more accessible the transcriptional machinery in aged tissues. Both local increases and decreased were identified throughout the mouse genome (Fig. 1C). Thus, our analyses suggest that the H3 nucleosomal landscape of aging mammalian tissues is subjected to localized remodeling, with both regions increased and decreased occupancy.

**Figure 1:**
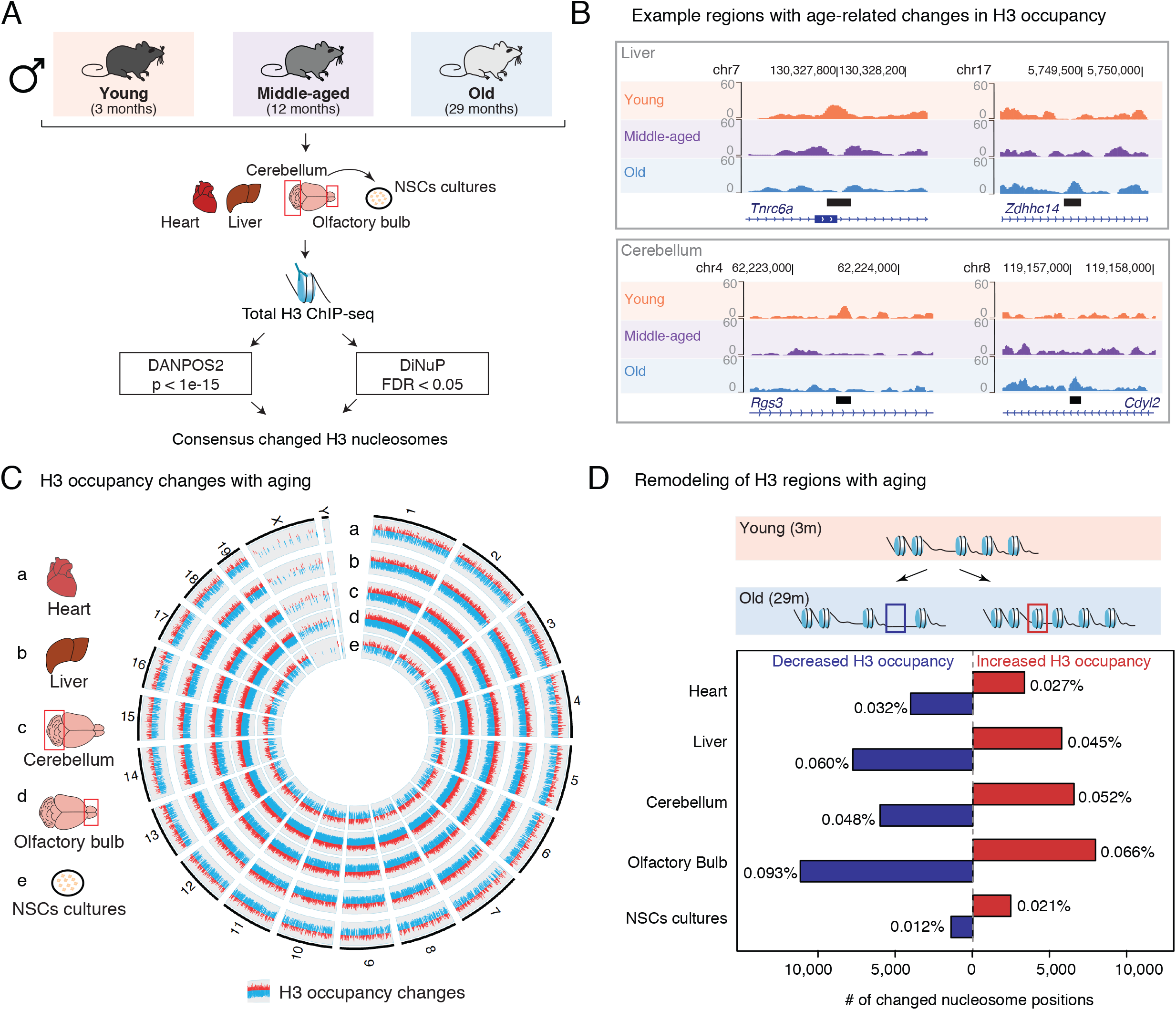
The genome-wide H3 nucleosomal landscape of mouse aging in four tissues and one cell types. (A) Experimental and analytical data setup. (B) UCSC Genome Browser Shots for examples of significantly remodeled nucleosomal loci in the liver (Top) or cerebellum (Bottom) samples. chr: chromosome. Coordinates are relative to the mm9 genome build. (C) Circular genome plot showing the genomic distribution of significantly remodeled H3 nucleosomes in Heart (a), Liver (b), Cerebellum (c), Olfactory Bulb (d) and primary NSC cultures (e). Note that there is no obvious clustering on specific chromosomes. (D) Barplot of frequencies of regions with increased (red) or decreased (blue) H3 occupancy with aging. The percentage of all detected nucleosomes with increased or decreased occupancy is reported next to the bar.

Because changes in global H3 levels may adversely affect our ability to properly call remodeled nucleosomal regions, we decided to assay potential changes in H3 protein levels with aging using western blot in 2 of the tissues for which we reanalyzed ChIP-seq datasets: cerebellum and liver from male mice (Fig. 2A). We obtained tissues from aging male C57BL/6N mice and extracted proteins for analysis. Our analysis of protein levels in these tissues is consistent with the absence of large-scale changes in protein expression levels of histone H3 with aging (Fig. 2B-C, **S2A-C**), and does not support the presence of consistent decrease in histone H3 expression in aging mammalian tissues. Importantly, we were able to recover the previously described increased levels of the histone variant macroH2A in aging mouse liver [23] (Fig. 2B), thus showing that we had the capacity to detect changes in histone protein levels. In addition, our analysis of the previously published RNA-seq dataset that accompanied the H3 ChIP-seq datasets [22] showed that there are no patterns of consistent transcriptional changes in histone H3 gene expression, with histone H3 transcripts both downregulated or upregulated with age (**Fig. S3**). Our observations are consistent with previous observations in an independent study of mouse liver aging [21]. Our western blot results are also consistent with lack of clear recurrent changes in the amount of spike-in ChIP-Rx reads from Drosophila S2 cells [24] recovered in the H3 ChIP-seq aging data [22] (**Fig. S4**). Thus, our analyses suggest that, in contrast to replicatively aged yeast [16] or senescent mammalian cells [17], core histone expression levels are relatively stable in chronologically aged mammalian tissues.

**Figure 2:**
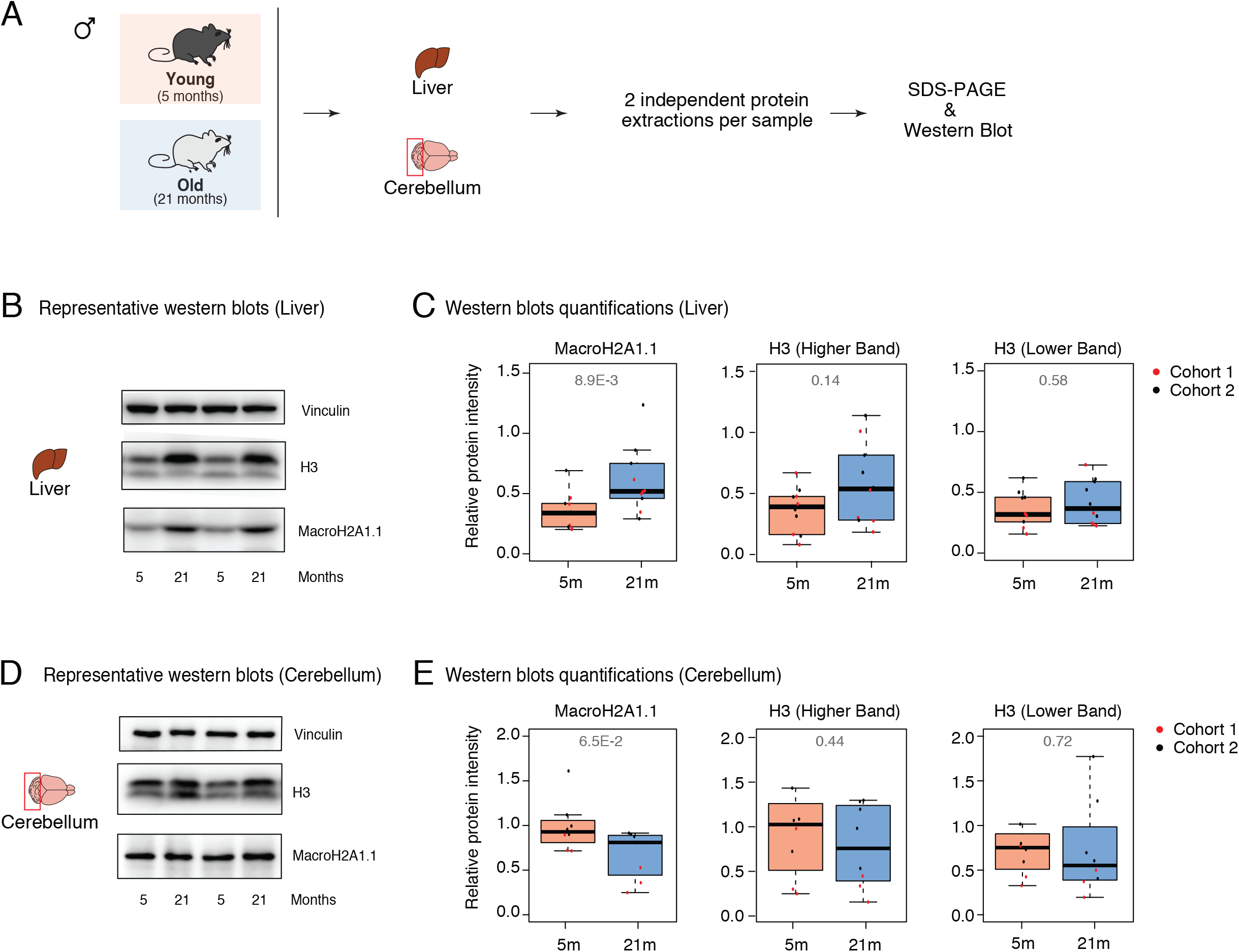
Analysis of H3 protein levels in aging mouse liver and cerebellum samples by Western blot. (A) Schematic illustrating methodology for protein quantification. (B,D) Representative western blot images for Vinculin (loading control), total H3, and macroH2A1.1 in liver (top) and cerebellum (bottom) protein extract. (C,E) Quantification of protein intensity from all Western blots from 5 vs. 21 months tissues corresponding (normalized to Vinculin loading control). The average relative protein intensity of technical replicates for each sample is reported. All quantified gels are provided in **Figure S2**. Exposure times for liver: vinculin (20 seconds), MacroH2A1.1 (50 seconds), H3 (2 seconds). Exposure times for cerebellum: vinculin (50 seconds), MacroH2A1.1 (50 seconds), H3 (4 seconds). P-values re reported on boxplots obtained with the non-parametric Mann-Whitney test.

### 2.2 Age-related changes in H3 occupancy are frequent at distal elements, but enriched in intronic regions

Significantly remodeled regions of H3 occupancy tended to occur most often at distal regions, 5 to 500 kb from annotated transcriptional start sites [TSS] (Fig. 3A-B). This observation is consistent with previous observations of age-related nucleosomal occupancy changes in aging livers using MNase-seq between 3 and 21 months of age [21], now also extending to heart, cerebellum, olfactory bulb and NSCs. In all tissue samples, when compared to the <u>genomic distribution of all detected nucleosomes</u> in our samples, sites in intronic regions were overrepresented compared to background nucleosomes (<36% of detected H3 nucleosomes *vs.* >39% of sites with remodeled H3 occupancy) (Fig. 3C-E). This was less pronounced in the NSCs samples, which were previously found to harbor much less age-related changes at the level of transcription and histone modifications [22]. This observation may indicate differential occupancy of regulatory elements (*e.g.* enhancers, silencers) in aging chromatin. In addition, we observed a general trend for regions with remodeled H3 occupancy to fall within regions close to genes (*i.e.* proximal promoter, coding exons, UTRs) more often than compared to background detected nucleosomes (Fig. 3C-E). This may suggest that lifelong patterns of transcriptional activity may ultimately lead to genomic remodeling of H3 occupancy.

**Figure 3:**
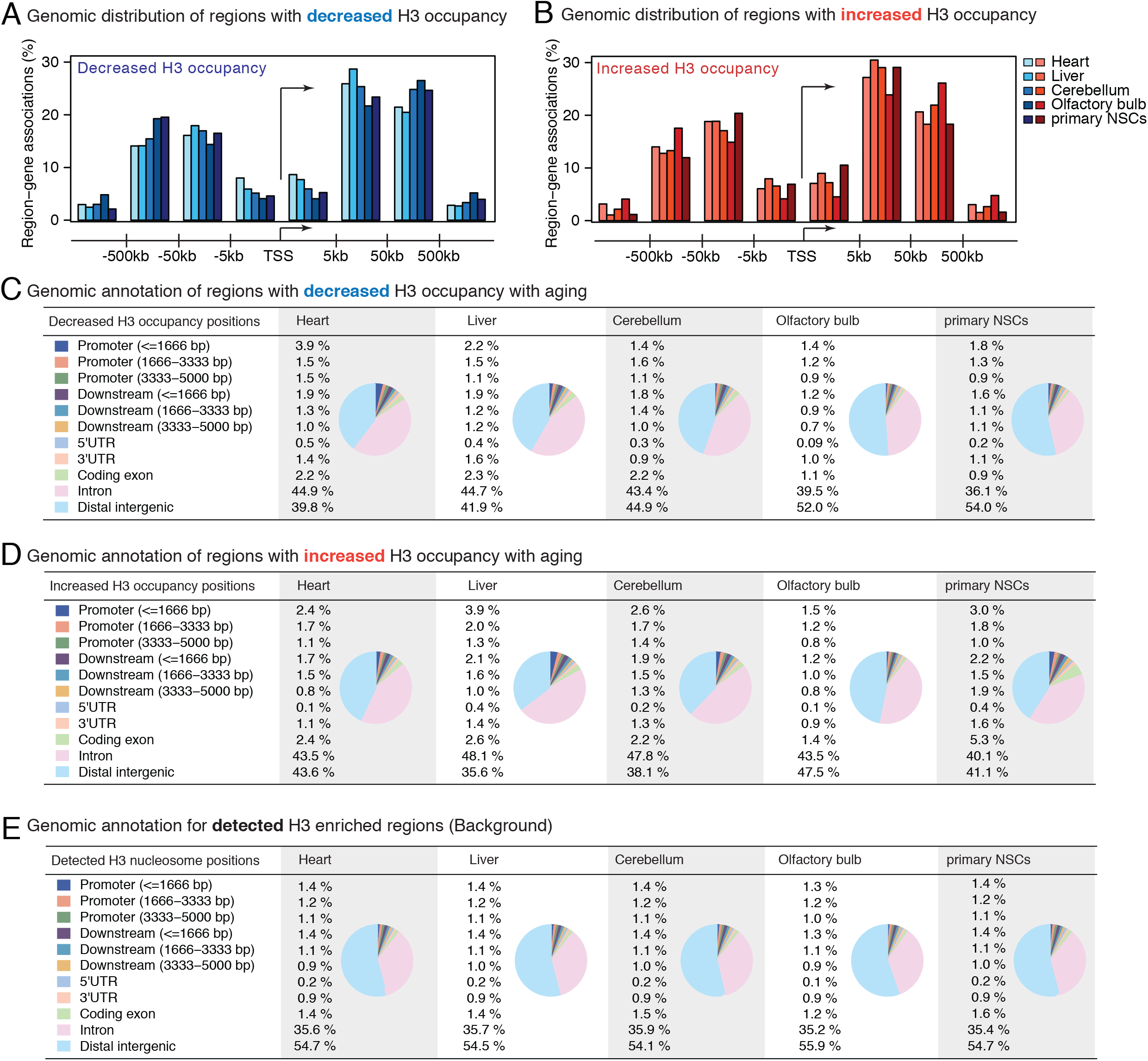
Genomic localization of age-remodeled nucleosomes. (A-B) Relative distance to annotated transcription start sites [TSSs] of nucleosomes with decreased (A) and increased (B) H3 occupancy during mouse aging. (C-E) Genome ontology analysis by CEAS of nucleosomes with decreased (C) or increased (D) H3 occupancy during mouse aging, compared to all detected (E) nucleosomes in the tissue.

Although informative, genome ontology analysis does not provide information on the function or activity of annotated elements. Chromatin-state annotation using combinations of chromatin modification patterns is a powerful approach to annotate tissue-specific activity patterns of regulatory regions [25, 26]. Thus, to evaluate which chromatin states are most affected by H3 occupancy remodeling, we decided to build tissue-specific chromHMM models by using available ChIP-seq data [25, 26] (**Table S1A-B**; **Figure S5A-G**). Interestingly, the majority of remodeled nucleosome outside of “low signal” regions resided in weak/poised enhancers (**Table S1A-B**), consistent with coordinated chromatin remodeling at regulatory regions during aging.

### 2.3 Differential enrichment of putative regulatory motifs is consistent with increased accessibility of pro-inflammatory gene programs

To understand the potential biological significance of remodeled regions of H3 occupancy in aging mouse tissues, we leveraged the GREAT annotation tools. We focused on the annotation “Molecular Signature Database motifs enriched at promoters” to identify potential associations to differential transcription factor behavior (Fig. 4A-B). The majority of motifs associated to changed regions that were recurrently enriched (3 or more datasets) were linked to Forkhead transcription factors [TFs] (*e.g.* FOXO4, FOF2, FOXL1, FOXA1, etc.), both at regions of increased and decreased H3 occupancy (Fig. 4). These observations are consistent with the previous MNase-seq analysis of aging liver [21], and show that remodeling of Forkhead TF regulated sites may be a general features of chromatin accessibility during aging. Interestingly, Forkhead TFs can act as ‘pioneer’ factors and directly remodel chromatin; for example, Foxo1 and Foxa2 are able to bind nucleosomal DNA, and lead to chromatin decompaction [27, 28]. FOXO factors are known to act as pro-longevity genes [29], and may thus help remodel relevant chromatin regions during the aging process. Interestingly, in an associated RNA-seq dataset, transcriptional targets of Forkhead TFs (*i.e.* FOXO1, FOXO3, FOXO4, FOXA1 and FOXA2) were found to be significantly transcriptionally upregulated with aging [22], consistent with the notion that Forkhead TF transcriptional target networks may be remodeled downstream of chromatin remodeling during aging.

**Figure 4:**
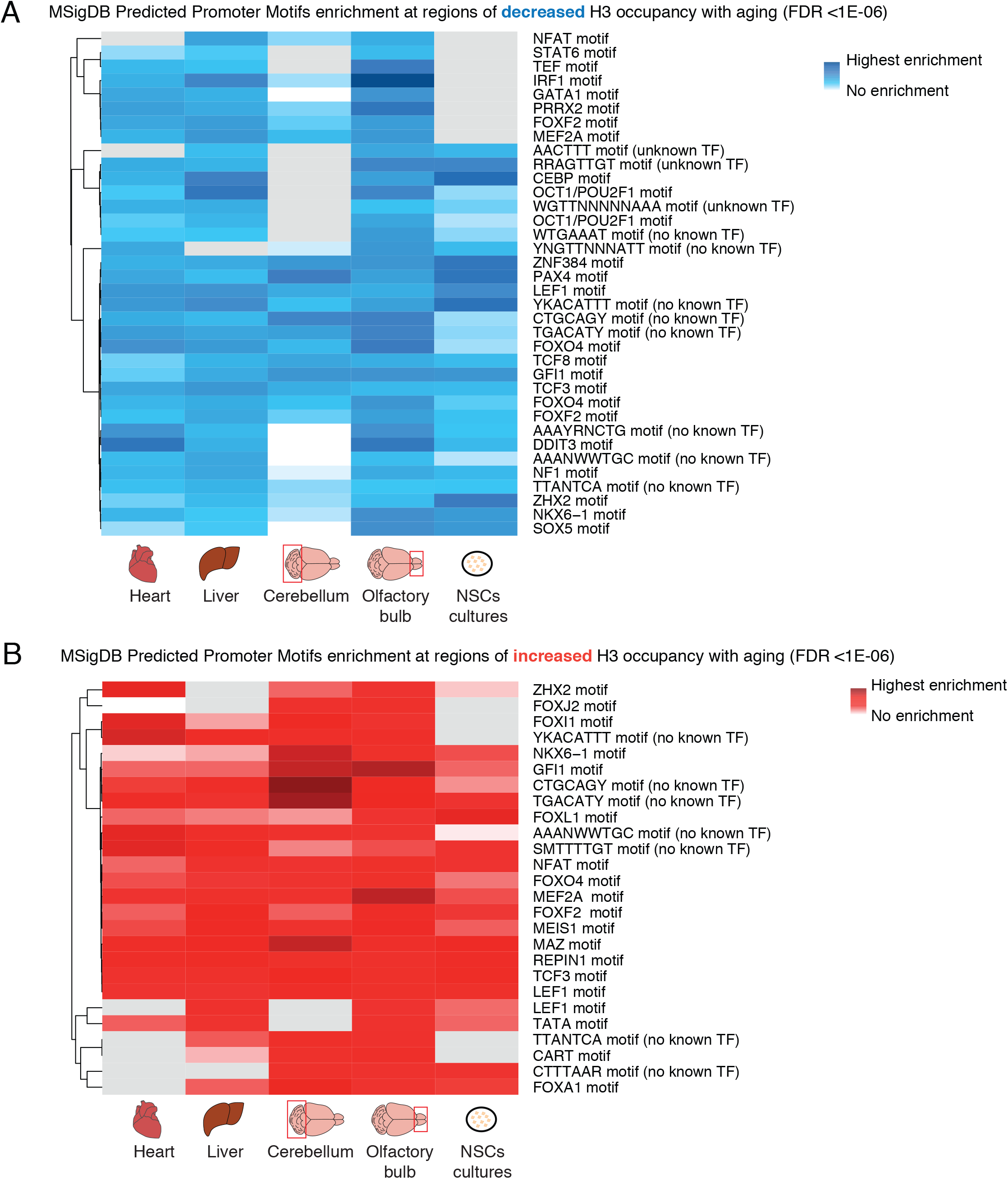
Enrichment of putative transcription factor targets for genes associated to age-remodeled nucleosomes. Remodeled nucleosomes with decreased (A) or increased (B) H3 occupancy with aging were given as inputs to the GREAT annotation portal. Results from the MSigDB transcription factor target annotation type are shown. Only annotations significant in 4 of the 5 tissues, with FDR <1E-06 are reported.

Consistent with the overall ‘omic’ signature of inflammatory response with age at the transcriptomic and histone modification levels [22], regions with decreased H3 occupancy (*i.e.* regions likely to be more accessible and active with aging) were recurrently enrichment for association to inflammation-related TF motifs (Fig. 4A), including motifs associated with IRF1, STAT6 and TCF3. Consistently, transcriptional targets of IRF8 and TCF3 were previously noted to be significantly upregulated with aging across tissues [22]. Thus, remodeling at the chromatin at the levels of nucleosomal occupancy and histone modifications may be partially coordinated by inflammatory TFs, and mechanistically underlie the induction of innate immune response pathways with aging.

## 3. Discussion

To understand the effect of aging on nucleosome occupancy with aging, we have reanalyzed transcriptomic and epigenomic maps in young, middle-aged, and old mice from a variety of tissues and cells known to show functional decline with aging (*i.e.* heart, liver, cerebellum, olfactory bulb and cultured primary NSCs) [22]. We also complemented the genomic study with lower-throughput western blot on total histone protein levels in aging tissues. To our knowledge, this study represents the most comprehensive analysis of histone expression and occupancy changes with mammalian aging to date. We hope that it will serve as a resource for the aging research community.

Based on our observations and previously published reports, it is likely that dramatic decreased nucleosome occupancy only occurs in a cell-type-or context-specific manner with aging and is not the rule in mammalian tissues. To note, it is also possible that MNase and ChIP-seq based methods may not be sensitive enough to detect subtle distribution changes, which may become detectable with higher-resolution methods. However, motifs enriched in regulatory regions of genes with remodeled nucleosomal occupancy revealed potential regulation by multiple inflammatory transcription factors, such as STAT6 and IRF8, which is consistent with transcriptional trends observed with aging [22]. Thus, changes in underlying accessibility of DNA to transcription factors may play a mechanistic role in the activation of inflammatory gene programs during mammalian aging.

In the long-term, understanding not only changes in histone post-translational modification patterns, but also changes in the underlying distribution of histones on the chromatin throughout life and in response to various stimuli will be key in understanding novel aspects of the “histone code” [4] regulation of biological processes.

## 4. Materials and Methods

### Mouse husbandry

All animals were treated and housed in accordance to the Guide for Care and Use of Laboratory Animals. All experimental procedures were approved by USC’s Institutional Animal Care and Use Committee (IACUC) and were in accordance with institutional and national guidelines. Male C57BL/6N mice at different ages were obtained from the National Institute on Aging (NIA) aging colony at Charles Rivers. Mice were acclimated at the animal facility at USC for 2-4 weeks before euthanasia. All animals were euthanized between 9-11am, and a variety of tissues were collected, snap frozen in liquid nitrogen, and stored at −80°C until further handling. No live animals were censored.

### Preparation of protein samples

Proteins were extracted from frozen liver and cerebellum samples. Each sample was split in two portions in order to carry out two independent protein extractions. At the times of protein extraction, tissues were maintained on ice, cut, and transferred to MP Biomedicals Lysing Matrix D tubes (Fisher Scientific Cat. No. MP116913050) containing 1X RIPA Lysis Buffer (Millipore Cat. No. 20-188) and 1X Halt Protease and Phosphatase Inhibitors (Thermo-Fisher Cat. No. 78440). Tissues were homogenized (1 cycle for cerebellums; 3 cycles for livers of 30 seconds at 3500 rpm), and lysates were centrifuged to remove debris and insoluble fraction at 10,000g and 4°C for 10 min. The supernatants were used as tissue protein lysates. Liver samples were centrifuged an additional time before collecting lysates in order to minimize fat transfer. Protein lysate concentrations were quantified using the Pierce BCA Protein Assay Kit (ThermoFisher Cat. No. 23227) following manufacturer’s instructions. Lysates were diluted to the same final protein concentration, mixed with Laemmli sample buffer (Biorad Cat. No. 1610737), and denatured by boiling at 95°C for 10 min.

### SDS-PAGE and Western Blot

For all cerebellum samples, 4–20% Mini-PROTEAN TGX Precast Protein Gels (Biorad Cat. No. 4561095) were used. For all liver samples, polyacrylamide gels were prepared with a 5% stacking layer (388 mM Tris pH 8.8, 4% Acrylamide:Bis 29:1, 1% SDS, 1% APS, 0.1% TEMED) and a 10% separating layer (388 mM Tris pH 8.8, 10% Acrylamide:Bis 29:1, 0.9% SDS, 0.9% APS, 0.04% TEMED) in Mini-PROTEAN cassettes (Biorad Cat. No. 4560005). Electrophoresis cells were filled with TAE Running Buffer (Bioland Cat. No. TAE02) before loading gels with 30 μg of protein per well. Precision Plus Protein Standards (Biorad Cat. No. 161-0374) were used to track protein mass. Gels were prepared in duplicates, one for transferring onto PVDF Blotting Membrane (Sigma Cat. No. 10600021), and one for total protein staining by Coomassie blue following manufacturer recommendations (Biorad Cat. No. 1610436, 1610438). Protein transfers were conducted in Transfer buffer (48 mM Tris base, 39 mM Glycine, 1.28 mM SDS, 20% ethanol) at 4°C for 1 hour with 250 mA of constant current. After transfer, membranes were blocked with 5% Bovine Serum Albumin [BSA] (Cat. No. AK8917-0100) in Tris-buffered saline (TBS, Cat. No. J62938) (liver) or TBS-10%Tween [TBST] (cerebellum) for 1 hour. All primary and secondary antibodies were prepared using 5% BSA in TBST. After blocking, membranes were cut into three pieces along the ~75 kD and ~25 kD molecular mass as indicated by protein standards, and incubated overnight at 4°C with Anti-Vinculin antibody (1:2000, Abcam ab91459), MacroH2A1.1 (D5F6N) antibody (1:1000; Cell Signaling Technologies 12455S), or Anti-Histone H3 antibody (1:1000 for liver and 1:5000 for cerebellum, Abcam ab1791), based on the expected molecular weights of the proteins to be assayed. Membranes were then washed with TBST three times and incubated with Goat Anti-Rabbit IgG H&L (HRP) secondary antibody 1:10,000 dilution (Abcam ab205718) for 1 hour at room temperature. Membranes were washed with TBST three times before incubating with SuperSignal West Pico PLUS Chemiluminescent Substrate (Thermofisher Cat. No. 34580) for 5 minutes. Protein signals were detected using an Azure c280 imaging system. Protein band quantifications were conducted using the gel analysis component of ImageJ and data was plotted using the R statistical software. For each sample, the average of 2-4 technical replicates are reported.

### ChIP-Seq data processing

H3 ChIP-seq data was obtained from the NCBI Short Read Archive (SRA) accession SRP057387 (BioProject PRJNA281127) and processed as described previously [22]. Briefly, sequencing reads were trimmed using Trimgalore v0.3.1 (https://github.com/FelixKrueger/TrimGalore) to retain high-quality bases with phred score >15 and a post-trimming length og 36bp. Reads were mapped to the genome using bowtie version 0.12.7. PCR duplicates were removed using FIXseq (fixseq-e2cc14f19764) [30].

### Nucleosome remodeling analysis

Total H3 ChIP-seq data was used to assess changes in nucleosome occupancy with aging. Nucleosome calls are derived from our previous study [22]. Briefly, since existing pipelines do not have time-series capability, we performed analyses comparing the young (3m) and old (29m) samples in each tissue type to assess the impact of aging. Differential occupancy position between 3 and 29 months were independently called using DANPOS (danpos v2.2.2) (p < 10^−15^) [31] and DiNup (dinup v1.3) (FDR < 5%) [32]. Only positions called as differential by both algorithms were considered as robustly changing with age and further analyzed.

### Analysis of genomic distribution of remodeled regions

Significant regions of interest (*i.e.* nucleosomes) were annotated to the gene with the closest transcription start site using HOMER [33]. We extracted and plotted the distance between the region of interest and closest annotated TSS. Basic genome ontology analysis was also performed with HOMER. The relative enrichment of remodeled regions compared to genomic elements was performed using the Cis-regulatory Element Annotation System [CEAS] software v1.0.2 [34].

### Analysis of histone gene expression in RNA-seq data

DEseq2 normalized RNA-seq expression count tables were derived from our previous study [22]. Official gene symbol for histone genes were obtained from BioMart, and genes encoding core or variants of histone H3 were selected for further analysis. Genes were considered detectable if at least 1 read was detected across all samples in a specific tissue. Expression for detectable histone H3 genes was then reported as a boxplot for each age, shown in **Fig. S3**.

### Analysis of chromatin states with chromHMM

To train chromHMM models [25, 26], we obtained ChIP-seq datasets (*i.e.* H3K4me3, H3K27ac, H3K4me1, H3K27me3, Pol2, CTCF and DNAse/ATAC) in the studied mouse tissues [35] and in cultured NSCs [22, 36, 37]. Mapped sequencing reads were fed to the chromHMM v1.12 algorithm to learn 10 states across the 5 conditions (*i.e.* heart, liver, cerebellum, olfactory bulb and NSCs). Chromatin state predictions were generated using ChromHMM v1.12 [25, 26]. The model was built using H3K4me3, H3K27ac, H3K4me1, H3K27me3, Pol2, CTCF and DNAse/ATAC from young adult samples (**Table S2** for dataset accession numbers accessions). Default parameters were used for training. For prediction of states, a 10-state model was empirically determined to most closely represent the expected states (*e.g.* promoter-like states found near TSSs). The final emission properties learned by ChromHMM are reported in **Fig. S5A-G**. For ease of interpretation, similar states were subsequently merged, resulting in 8 summarized main states: low signal, active enhancer, weak/poised enhancer, inactive/poised promoter, polycomb repressed, insulator, active promoter, and flanking promoter.

### Functional enrichment analysis of remodeled regions using GREAT

Functional term enrichment analysis was conducted by comparing differentially enriched H3 regions to all detected H3 regions using GREAT v3.0.0 [38]. Genomic coordinates of peaks (in the form of bed files) were used. For ease of graphical representation, terms enriched in at least 3 of the 5 conditions at FDR < 10^−6^ were selected for analysis and plotting.

### Code Availability

All new analytical code will be made available on the Benayoun laboratory Github repository.

## Acknowledgements

J.B. was supported by NIA T32AG05237 and NSF graduate research fellowship DGE-1842487. B.A.B is supported by NIA R00AG049934, an innovator grant from the Rose Hills foundation, a seed grant from the NAVIGAGE foundation, and a generous gift from the Hanson-Thorell Family.

## Conflict of interest

The authors declare that they have no conflict of interest to disclose.

## Supplementary Figure Legends

**Figure S1: The genome-wide H3 nucleosomal landscape of mouse aging in four tissues and one cell types.** (A) Heatmap of H3 ChIP signal centered on regions called as increased or decreased with aging. Note the increased or decreased signal at these sites.

**Figure S2: Analysis of H3 protein levels in aging mouse liver and cerebellum samples by Western blot (continued).** (A) Western blots for Vinculin, H3, and macroH2A1.1 from livers of 2 cohorts of male aged mice (5 months and 21 months). Samples were run on a 10% SDS-PAGE gel, with extractions 1 and 2 denoting technical duplicates derived by cutting each tissue in two pieces. Homogeneous protein loading was also assessed by Coomassie blue staining. Exposure times: vinculin (50 seconds), macroH2A1.1 (50 seconds), and H3 (4 seconds). (B) Western blots for Vinculin, H3, and macroH2A1.1 from cerebellum of 2 cohorts of male aged mice. The first cohort had only animals with 5 and 21 months of age, and the second cohort had 5, 12, 21, and 25 month old mice. Samples were run on a 4-20% SDS-PAGE gradient gel. Exposure times: vinculin (8 minutes), macroH2A1.1 (8 minutes), and H3 (3 seconds). (C) H3 blots (3 second exposure) for cerebellum and liver samples run on a 15% polyacrylamide gel. Apparent molecular weight differences for the H3 bands between cerebellum and liver samples in A and B appear to be due to differences in the percentages of the separating gels.

**Figure S3: Analysis of H3 protein levels in aging mouse liver and cerebellum samples by Western blot (continued).** (A-E) Expression by RNA-seq of H3-encoding genes in heart (A), liver (B), cerebellum (C), olfactory bulb (D) and primary NSC cultures (E). Note that there is no general rule as to increased or decreased transcript levels for H3-encoding genes in these tissues.

**Figure S4: Analysis of H3 ChIP-sea reads mapping to mouse vs. drosophila genome.** Reads were mapped to the mouse mm9 build or the drosophila dm3 build to determine the reads from the mouse aging tissue (mm9) and the reads from the spiked-in S2 cells [22]. If no change in H3 levels occur on chromatin, all values will be at 1. If there is a decreased in H3 loaded onto chromatin with aging, values will reliably drop below 1.

**Figure S5: ChromHMM model parameters.** (A) Emission parameters of the trained model, and predicted function based on enriched histone marks. (B) Transition parameters of the trained model. (C-G) Enrichment of various genomic features for the different learned states.

## Notes

https://github.com/BenayounLaboratory/Nucleosome_aging_2019

