## Supplementary figures and images for "Remodeling of the H3 nucleosomal landscape during mouse aging"

### Fig S1

Figure S1

A H3 ChIP-seq signal at significantly remodeled regions

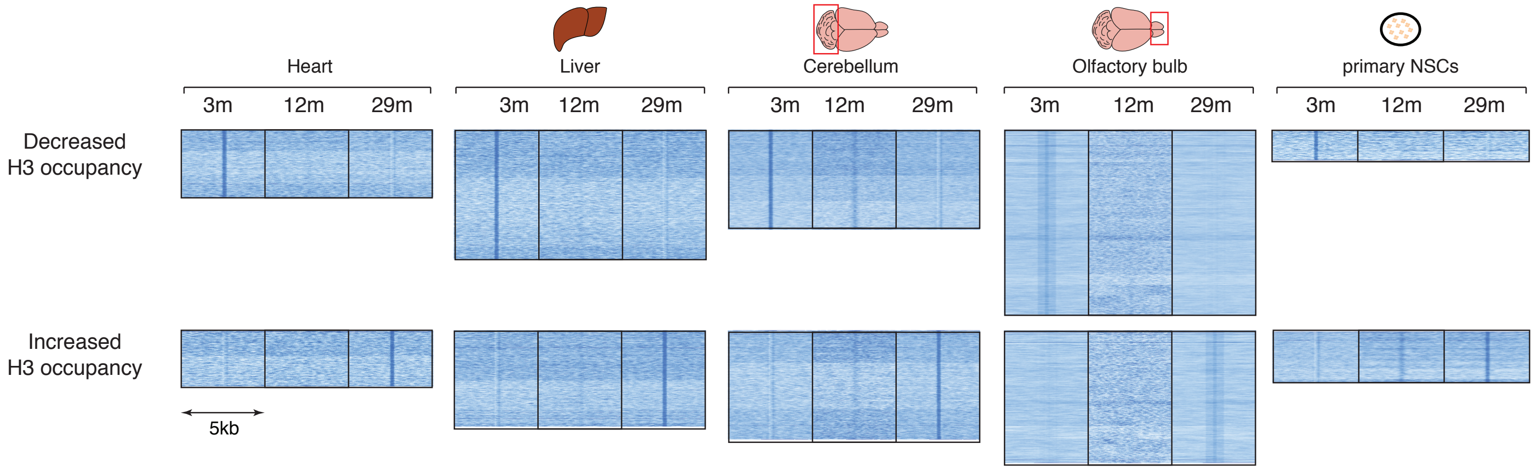

### Fig S3

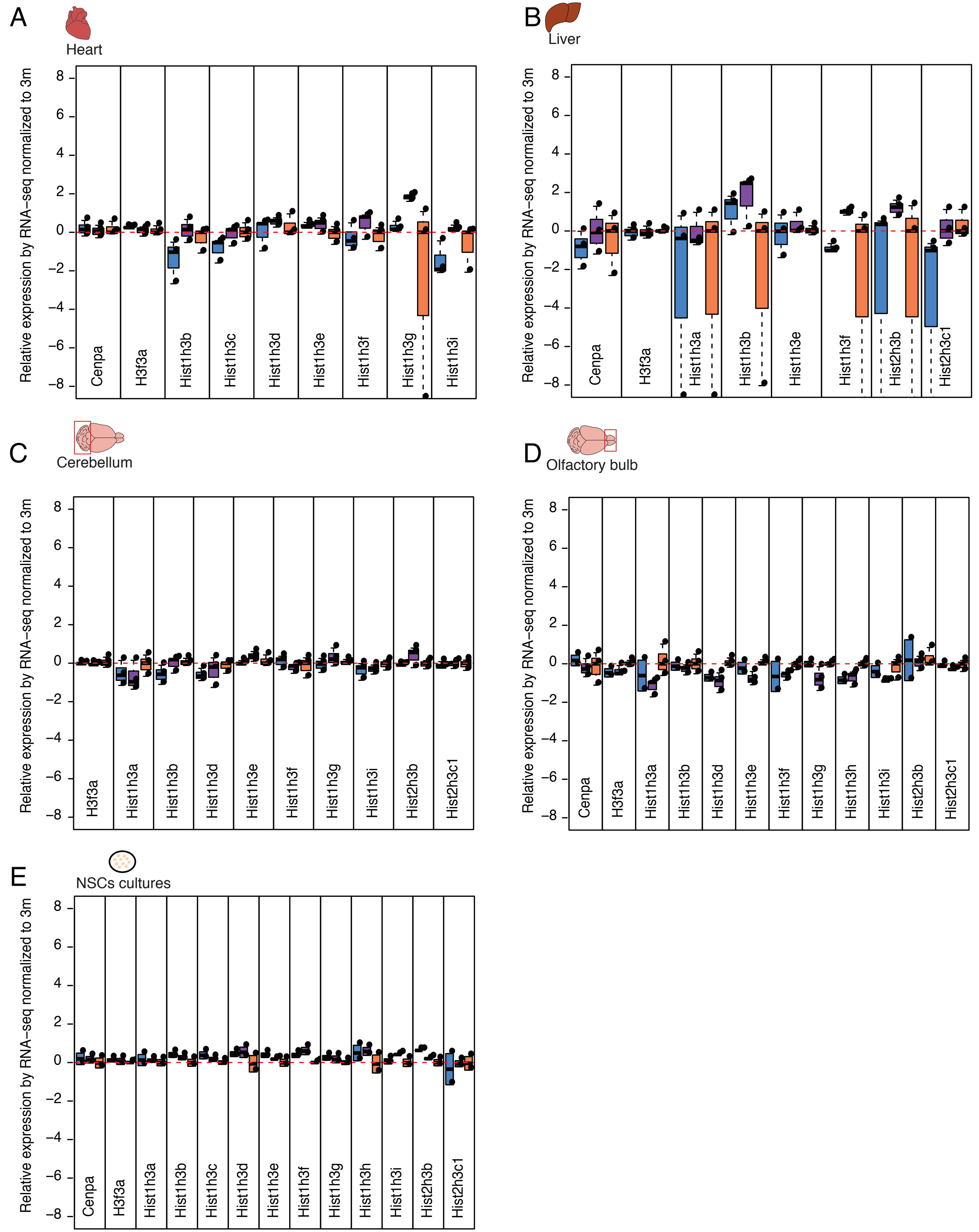

### Fig S4

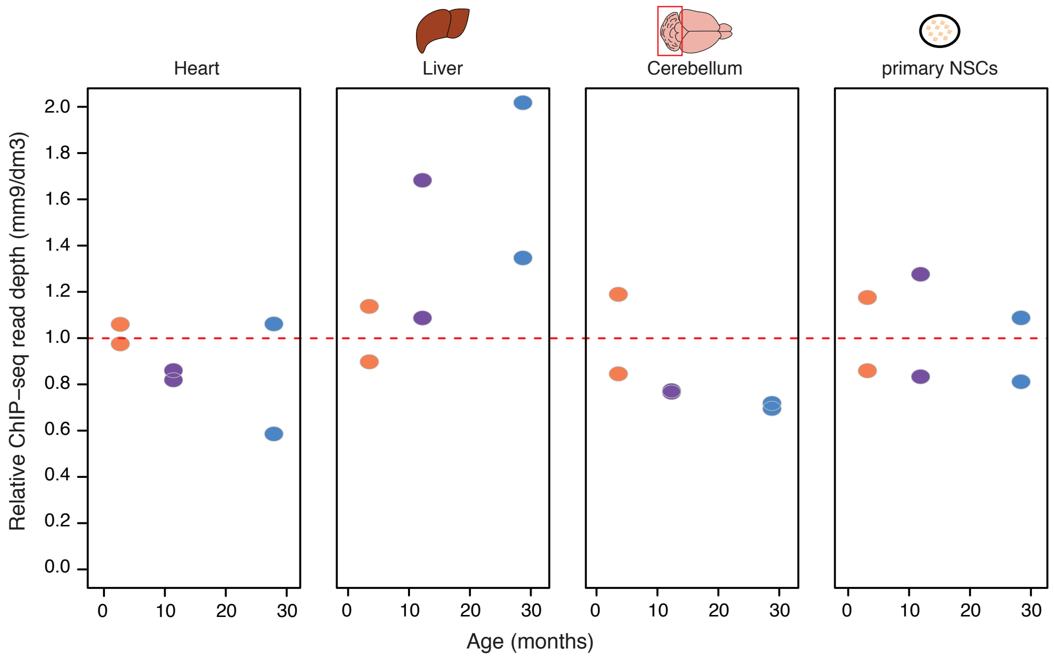

### Fig S5

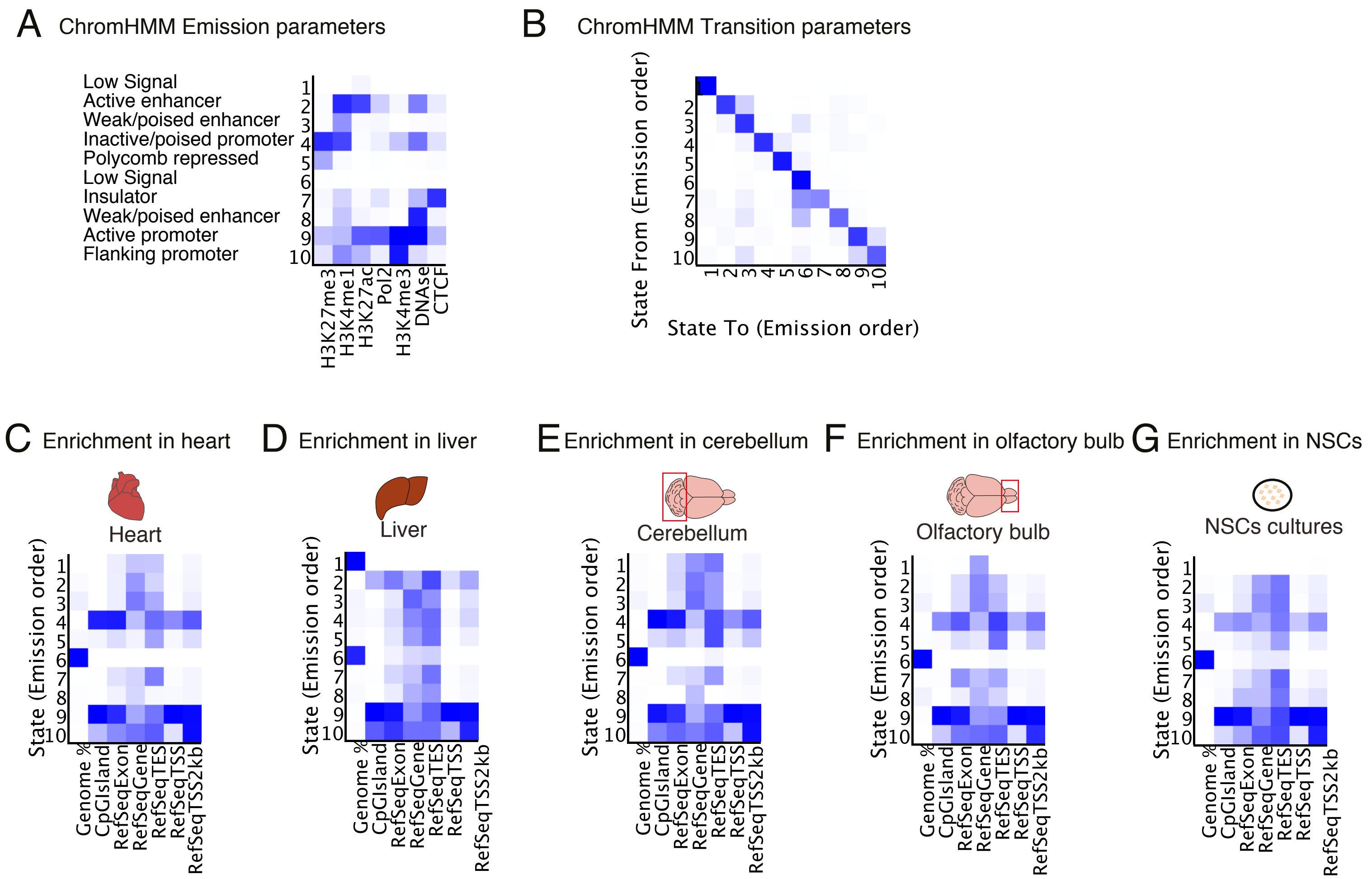
