## Supplementary material for "Remodeling of the H3 nucleosomal landscape during mouse aging": Fig S2

Figure S2. Western blot images for aging mouse liver and cerebellum

A Western blot images from liver tissues, run on 10% polyacrylamide gels

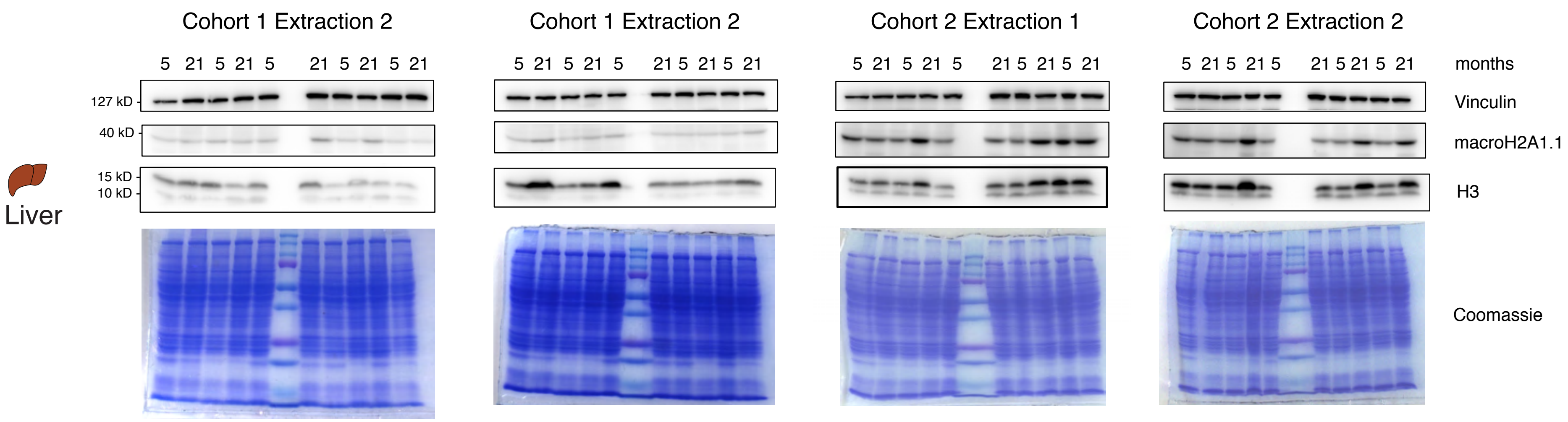


B Western blot images from cerebellum tissues, run on 4–20% Mini-PROTEAN TGX Precast Protein Gels

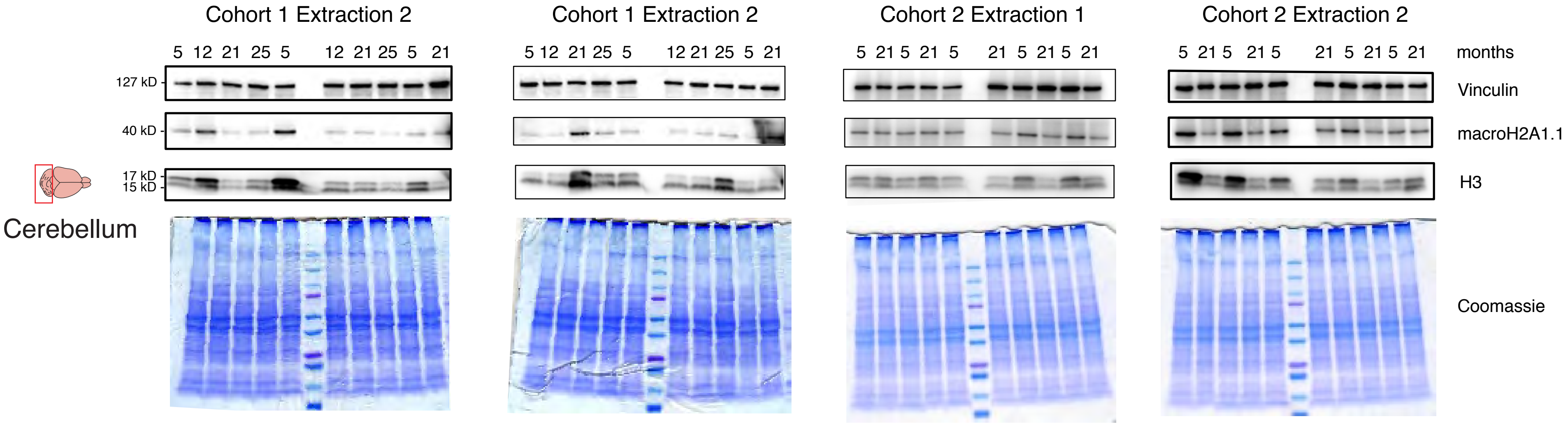


C Western blot images from liver and cerebellum tissues, run on a 15% polyacrylamide gel

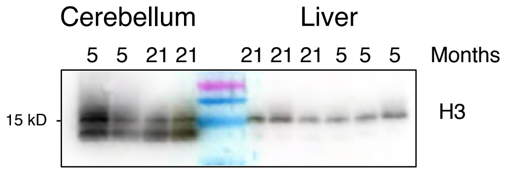
